# Immune function of the serosa in hemimetabolous insect eggs

**DOI:** 10.1101/2021.09.27.462031

**Authors:** Chris G.C. Jacobs, Remy van der Hulst, Yen-Ta Chen, Ryan P. Williamson, Siegfried Roth, Maurijn van der Zee

**Author notes:** Corresponding author:* Maurijn van der Zee, Institute of Biology, Leiden University, Sylviusweg 72, 2333 BE Leiden, The Netherlands. Link to deposited data: https://www.ncbi.nlm.nih.gov/geo/query/acc.cgi?acc=GSE100429.

## Abstract

Insects comprise more than a million species and many authors have attempted to explain this success by evolutionary innovations. A much overlooked evolutionary novelty of insects is the serosa, an extraembryonic epithelium around the yolk and embryo. We have shown previously that this epithelium provides innate immune protection to eggs of the beetle *Tribolium castaneum*. It remained elusive, however, whether this immune competence evolved in the *Tribolium* lineage or is ancestral to all insects. Here, we expand our studies to two hemimetabolous insects, the bug *Oncopeltus fasciatus* and the swarming grasshopper *Locusta migratoria*. For *Oncopeltus*, RNA sequencing reveals an extensive response upon infection, including the massive upregulation of antimicrobial peptides (AMPs). We demonstrate antimicrobial activity of these peptides using *in vitro* bacterial growth assays, and describe two novel AMP families called Serosins and Ovicins. For both insects, qPCRs show immune competence of the eggs when the serosa is present, and in situ hybridizations demonstrate that immune gene expression is localized in the serosa. This first evidence from hemimetabolous insect eggs suggests that immune competence is an ancestral property of the serosa. The evolutionary origin of the serosa with its immune function might have facilitated the spectacular radiation of the insects.

## 1. Introduction

Insects are extraordinarily successful and radiated into more than a million species [1]. Over the last century, several authors have attempted to explain the success of the insects by the origin of key innovations such as wings or metamorphosis [2, 3]. A much overlooked evolutionary novelty, however, is the serosa, an extraembryonic epithelium in insect eggs that covers the yolk and embryo [4–7]. This epithelium is a synapomorphy of the insects and is not present in other arthropods such as crustaceans [4, 5, 7, 8]. The serosa secretes a cuticle that protects the embryo against desiccation [8–11]. Thus, the origin of the serosa might have opened up a whole new range of terrestrial oviposition sites, facilitating the radiation of insects on land [7, 8].

Strikingly, the serosa was secondarily lost in a group of derived flies (the Schizophora) to which the well-studied model insect *Drosophila melanogaster* belongs [12, 13]. This makes the serosa a poorly studied epithelium by evolutionary and developmental biologists. In line with this absence, *Drosophila* eggs have extremely limited resistance to desiccation [14]. Desiccation is not the only challenge for insect eggs; they are also constantly challenged by pathogens [15–20]. The innate immune response of *Drosophila* eggs, is limited too [21]. It is not until stage 15 (one of the latest developmental stages when the ectoderm and trachea differentiate) that *Drosophila* eggs show upregulation of antimicrobial peptides after challenge [22]. Younger eggs cannot contain an infection of non-pathogenic bacteria [22].

In contrast to *Drosophila*, eggs of the mealworm beetle *Tenebrio molitor* can mount an extensive innate immune response, including upregulation of peptidoglycan recognition proteins (PGRPs), Toll, and AMPs [23]. These eggs do possess a serosa. Interestingly, inducible antimicrobial peptide expression was also found in the yolk-and-serosa fraction from *Manduca sexta* eggs, but not in isolated germ bands [24]. We have previously demonstrated that it is the serosal epithelium that harbors the innate immune response in eggs of the beetle *Tribolium castaneum* [25]. Serosal cells express mRNAs of antimicrobial peptides, serosa-less eggs do not upregulate immune genes, and bacteria propagate twice as fast in serosa-less eggs [25].

It remains unclear, however, whether immune competence of the serosa is ancestral to all insects, or evolved in butterflies and (tenebrionid) beetles. This is particularly elusive, as not only do the serosa-less *Drosophila* eggs lack an immune response, but so do the serosa-possessing eggs of the burying beetle *Nicrophorus vespilloides* [17]. In order to address this issue, we expand our studies to the milkweed bug *Oncopeltus fasciatus*, and the orthopteran *Locusta migratoria*, both belonging to the Hemimetabola, the basal main group of insects that show incomplete metamorphosis. Upon immune challenge by a Gram positive and Gram negative bacterium simultaneously, RNAsequencing reveals an extensive inducible immune response in eggs of the milkweed bug *Oncopeltus fasciatus*. We demonstrate the antimicrobial activity of two novel families of AMPs: Serosins and Ovicins. For both the bug and the locust, qPCRs show that eggs are immune responsive when the serosa is present. Using in situ hybridization, we find that transcripts of upregulated AMPs are located in the serosal cells but not in the young embryo of both insects. Our data provide evidence that immune competence is an ancestral feature of the serosa, and we discuss these data in an evolutionary perspective.

## 2. Materials and methods

### (a) Insect cultures

*Oncopeltus fasciatus* were supplied with sunflower seeds and water, and were kept under a 12h:12h light:dark cycle at 25 °C and 65% relative humidity (RH). Cotton wool was provided for egg lay. *Locusta migratoria* were ordered from https://www.sprinkhanenwinkel.com/ and kept in cages under a 13h:11h light:dark cycle at 32°C and 50% RH. Fresh grass was collected and fed to them daily. Pots with 60% peat and 40% sand were wetted until the mixture was sticky and provided for egg lay.

### (b) Infection and RNA isolation

Infection was performed using our previously described standardized infection method [25]. *Escherichia coli* (DMSZ 10514) and *Micrococcus luteus* (DMSZ 20030) were grown overnight in liquid LB at 37 °C. OD600 was measured and equal numbers of bacteria were spun down for 10 minutes at 4000g, and the pellets were mixed. *Oncopeltus* eggs of developmental stages A, B or C (see (e) qPCR) or 72-84h old *Locusta* eggs from at least three different egg cases, were then either left naïve, were pricked with a sterile tungsten needle (sterile injury), or were pricked with a tungsten needle that was dipped in this concentrated mixture of bacteria (septic injury). *Oncopeltus* eggs were kept at 25 °C; *Locusta* eggs at 32 °C. Six hours later, RNA was isolated from around 50-100 *Oncopeltus* eggs, or 6-9 *Locusta* eggs per treatment using Trizol (Invitrogen) extraction, followed by DNA digestion and column purification (Qiagen RNeasy). RNA quality was confirmed spectrophotometrically and by agarose gel electrophoresis.

### (c) *Oncopeltus* RNA sequencing analysis and genome annotation

Reads were checked for quality using FastQC v0.11.5 (http://www.bioinformatics.babraham.ac.uk) and trimmed using Trimmomatic (v0.36; HEADCROP:10 MINLEN:50 TRAILING:20). We assembled the trimmed paired end RNAseq reads from all samples using Trinity (v 2.4.0) applying the standard settings [26]. Then, we searched for immune genes in the genome [27] using BLAST, and checked gene predictions against our *de novo* assembly. In total, we modified 47 gene models of Official Gene Set v1.2 [28], merged 6 models into existing ones, and added 3 completely missing models. The full coding sequences of all identified immune genes can be found in the electronic supplementary material, table 1. Next, we mapped all reads to this updated version of the Official Gene Set and genome using hisat (v2.0.5) [29, 30]. Finally, the counts per gene were calculated using featureCounts (v1.5.2), and the data were analyzed using DEseq2 [31, 32]. A gff and fasta file of all updated gene models, the RNA sequencing data, and details of our expression analysis (R Markdown document) have been deposited in the NCBI’s Gene Expression Omnibus with accession number GSE100429 (https://www.ncbi.nlm.nih.gov/geo/query/acc.cgi?acc=GSE100429) [33].

**Table 1.** Infection induces an immune response in 48-64h *Oncopeltus* eggs. Fold upregulation of immune genes after sterile or septic injury based on whole transcriptome analysis in three biological replicates.OFAS numbers refer to *Oncopeltus fasciatus* Official Gene Set 1.2 [28]. Colours indicate amplitude of upregulation. White indicates <1.3 times or insignificant (p>0.05) upregulation. Asterisks indicate level of significance. Potential AMPs were predicted using the prediction software on the APD2 web site [42]. Upregulated genes after septic injury consist of 18 (potential) immune response execution genes, and 6 genes involved in the sensory machinery.

| category | OFAS | description | fold upregulation<br>upon sterile injury | fold upregulation<br>upon septic injury |
| --- | --- | --- | --- | --- |
| AMPs/effectors | 013883 | potential AMP Serosin-1 | 1.0 | 3173.8*** |
|  | 013884 | potential AMP Serosin-2 | 7.2 | 2606.2*** |
|  | 013885 | potential AMP Serosin-3 | 1.9 | 3994.9*** |
|  | 009139 | potential AMP Ovicin-1 | 1.8 | 1366.4*** |
|  | 009140 | potential AMP Ovicin-2 | 3.6 | 1595.7*** |
|  | 005103 | Defensin-1 | 8.2 | 2035.7*** |
|  | 005104 | Defensin-2 | 1.0 | 160.0*** |
|  | 005105 | Defensin-3 | 2.5 | 1136.6*** |
|  | 012870 | Oncocin | 0.7 | 108.9*** |
|  | 009533 | potential AMP | 0.4 | 40.3*** |
|  | 006190 | potential AMP | 14.1*** | 14.7*** |
|  | 011944 | matrix metalloproteinase | 14.5*** | 9.7*** |
|  | 011945 | matrix metalloproteinase | 14.0*** | 9.6*** |
|  | 011306 | laccase | 5.8*** | 11.9*** |
|  | 013043 | potential AMP | 4.8** | 5.3** |
|  | 009172 | potential AMP | 4.1*** | 3.7*** |
|  | 004519 | potential AMP | 4.8*** | 3.5*** |
|  | 018039 | potential AMP | 2.6*** | 2.3*** |
| sensory | 006421 | Serine protease | 3.5*** | 3.5*** |
|  | 009025 | Serine protease | 3.4*** | 4.0*** |
|  | 009027 | Serine protease | 3.1*** | 3.3*** |
|  | 006419 | Serine protease | 1.2 | 1.6** |
|  | 001578 | PGRP | 3.1** | 3.4*** |
|  | 011521 | Toll1 | 1.3* | 1.3* |

### (d) *Locusta* genome annotation and *Locusta* qPCR primer design

We searched the available *Locusta migratoria* genome https://i5k.nal.usda.gov/data/Arthropoda/locmig-%28Locusta_migratoria%29/ for *dorsal* and potential antimicrobial peptide sequences using local BLAST [34]. We identified 4 potential thamautins and 8 lysozymes, but could not design qPCR primers for them as sequences were very incomplete. Furthermore, we found dorsal, 3 Locustins, 9 Attacins, 6 Defensins, and 4 putative defense proteins (that we called “PDPs”) showing similarity to *Hyphantria cunea* Hdd11 [35]. We designed qPCR primers against all of these genes, but detected a reliable qPCR signal only for Locustin1, Attacin3, Attacin7, Defensin1, PDP1-3, and Dorsal. The sequences of used primers can be found in the electronic supplementary material, table 2.

### (e) qPCR

*Oncopeltus* qPCR primers were designed by Primer3 [36] using genome [27] and our transcriptome, and can be found in the electronic supplementary material, table 2. *Oncopeltus* cDNA was synthesized using the Cloned AMV First Strand Synthesis kit (Invitrogen); *Locusta* cDNA was synthesized using the cDNA Synthesis Kit (Promega). 2.5 ng of cDNA was used in each qPCR reaction. For *Oncopeltus*, Real-Time detection was carried out on a CFX96 Thermocycler (Biorad) using the SYBR Green I qPCR core kit (Eurogentec). Thermal conditions were 50°C for 2 minutes, 95°C for 10 minutes, 50 cycles of 95°C for 15 seconds, 60°C for 30 seconds and 72°C for 30 seconds, followed by a ramp from 65 to 95°C to confirm a single PCR product in a melting curve analysis. In total, qPCR was performed on 27 samples: three treatments (naïve, sterile injury, septic injury) were performed at three stages (stage A, 16-24h old eggs; stage B, 48-64h eggs; and stage C, 112-120h old eggs) in three biological replicates. Each qPCR was done in two technical replicates. From four potential reference genes (Ribosomal Protein L13a, Ribosomal protein 49, actin5cb, and heat shock protein Hsp90), RPL13a showed the most stable expression levels and was selected as internal control to calculate ΔC_T_ values. ΔΔC_T_ values after sterile or septic injury were calculated using the naïve egg samples as calibrator and fold upregulation was then calculated using the 2^-ΔΔC_T_^ method [37].

For *Locusta*, qPCRs were performed using the SsoAdvanced Universal SYBR Green Supermix (BioRad) at 95°C for 30 seconds, 40 cycles of 95°C for 15 seconds, 60°C for 30 seconds and 72°C for 30 seconds, followed by a ramp from 65 to 95°C for melting curve analysis (reads every 0.5°C) on six biological replicates, each analyzed in two technical replicates. From two potential reference genes (RPL13a and RPL32), RPL32 showed most stable expression, and was selected as internal control to calculate ΔC_T_. Upregulation after sterile or septic injury was then calculated using the 2^-ΔΔC_T_^ method with the naïve egg samples as calibrator [37].

### (f) In situ hybridization

For *Oncopeltus*, an approximately 350bp fragment of the AMP genes was amplified from cDNA of infected eggs using the primers

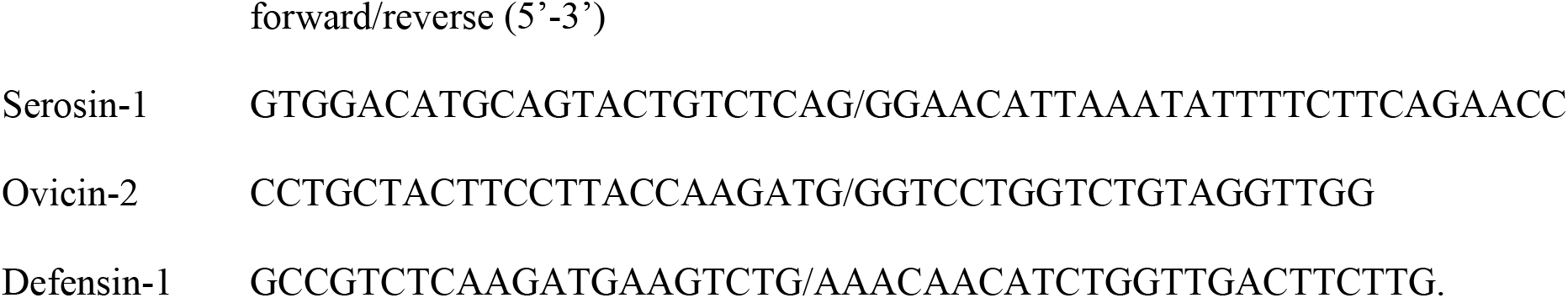

For *Locusta*, a 600bp fragment of dorsal and a 300bp fragment of Locustin were amplified from cDNA of infected eggs using the primers

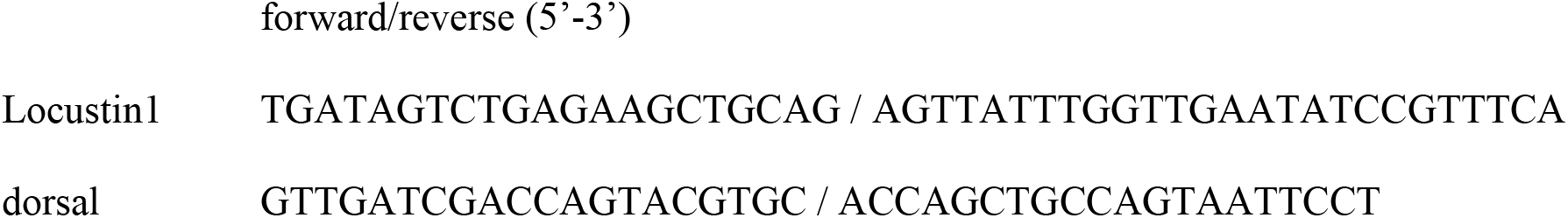

All fragments were cloned into the TOPO II vector (Invitrogen). Templates for *Locusta* probes were generated from positive clones (verified by Sanger sequencing) using the M13 primers. Plasmids containing *Oncopeltus* inserts (verified by analytical digest with PstI or HindIII), were linearized using XhoI (NEB) to generate a template for *in vitro* transcription. DIG-labelled in situ hybridization probes were transcribed *in vitro* using the SP6 or T7 MEGAscript kit (Ambion) with Roche RNA labelling mix (Roche). For *Oncopeltus*, embryo fixation (6h after infection) and in situ hybridization were performed as described by Liu and Kaufman [38]. For *Locusta*, 72-84h old infected or naïve *Locusta* eggs were fixed by perforating them 5-15 times with a tungsten needle and incubating them overnight in 9.75% formaldehyde in PBS, as suggested for *Schistocerca* [39]. Eggs were manually dissected to remove the yolk, often separating the embryo from the serosa. In situ hybridization was essentially performed as described for *Schistocerca* starting with a refixation [40], but after hybridization, unbound probe was washed away consecutively for 5, 10, 15 and 30 minutes, 4 times 1 hour, and finally overnight with posthybridization buffer at 55°C [40]. Sheep anti-dioxigenin-AP antibody (Roche) was used 1:5000 in blocking solution [40].

### (g) Bacterial growth assays

Predicted mature *Oncopeltus* peptides (see the electronic supplementary material, table 3) were ordered from Biomatik (Cambridge, Ontario, Canada). *Escherichia coli* (DMSZ 10514) and *Micrococcus luteus* (DMSZ 20030) were diluted to 1*10^6^ CFUs/ml in PBS. Agar plates were made by mixing 1 ml of this suspension in 10 ml of liquid LB agar. Holes were punched in these agar plates using a sterile cork bore. 10ul of a 100 μM peptide solution, an Ampicillin solution (1 mg/ml against *E. coli*; 100μg/ml against *M. luteus*), or sterile water was pipetted in such a hole. Plates were photographed after an overnight incubation at 37 °C. The experiment was carried out in triplicate.

## 3. Results

To assess a potential transcriptional response of *Oncopeltus fasciatus* eggs upon infection, we compared the complete transcriptomes of 48-64 h old eggs that were either left naïve, were sterilely injured, or were pricked with a needle dipped in a mix of *Escherichia coli* and *Micrococcus luteus* bacteria (septic injury). RNA sequencing of three biological replicates per treatment yielded in total more than 210 million reads of which more than 80% could be mapped to the *Oncopeltus fasciatus* genome [41]. Infection induced significant upregulation (>1.3 fold upregulation, adjusted p-value <0.05) of 67 genes, of which 48 were also significantly upregulated upon sterile injury (see the supplementary material, tables 4 and 5). Twenty-four of the upregulated genes, listed in table 1, were identified as immune genes based on similarity to *Drosophila melanogaster* immune genes or by potential antimicrobial properties (see below, [42]). Other upregulated genes include four ankyrin repeat containing proteins, two spinster-like proteins, a takeout-like protein, and 13 genes of unknown function or without clear homology to *Drosophila* genes (see the electronic supplementary material, table 4). The upregulated immune genes comprise genes involved in sensing infection, such as serine proteases; genes involved in signaling, such as the Toll receptor and a PGRP, but mainly include effector genes like potential AMPs, metalloproteinases and laccase (table 1).

A heatmap visualizing the number of reads of all immune genes identified in the genome reveals that some immune genes are constitutively expressed at high levels, such as the recognition genes GNBP (Gram-negative binding protein) and C-type lectins, and components of the JNK, IMD and Toll signaling pathways (see the electronic supplementary material, table 6). Interestingly, it is *cactus-1* that is expressed at high levels rather than *cactus-3* which has been shown to be involved in early dorsoventral patterning [43]. This suggests that subfunctionalization may have taken place among the *Oncopeltus cactus* paralogs. Upregulation upon infection was mostly detected among the execution genes (see the electronic supplementary material, table 6). Taken together, our RNA sequencing analysis indicates that 48-64h old *Oncopeltus* eggs can mount an innate immune response.

Some of the upregulated effector genes (table 1) were straightforward to identify as AMPs, such as OFAS005103, OFAS005104 and OFAS005105 that bear similarity to described Defensins. The OFAS009139 and OFAS009140 sequences, however, did not produce any BLAST hits but in *Halyomorpha halys*. We call them Ovicin1 and 2, respectively. OFAS013883-OFAS13885 only had similarity to undescribed Hemipteran peptides (in *Cimex lectularius, Gerris buenoi, Halyomorpha halys, Lygus hesperus, Rhodnius prolixus* and *Triatoma infestans*). We call them Serosin1-3. All these peptides were predicted to be antimicrobial based on their size, charge, hydrophobicity, and Proline- or Glycine-content [42] (figure 1*a*). To verify their antimicrobial activity, and to verify the functionality of the detected response in general, we synthesized the predicted mature peptides (supplementary table 3) and tested their effect on bacterial growth *in vitro*. 10 μl of a 100μM peptide solution was pipetted into wells that were punched in an LB agar plate containing 1×10^5^ CFUs/ml of *E. coli* or *M. luteus*. After an overnight incubation at 37 °C, no zones of inhibitions were observed around wells with Serosin-1 or Ovicin-2, but clear zones of inhibition demonstrated that Serosin-2 and Defensin-1 are active against *E. coli*, and that Ovicin-1 and Serosin-3 are active against *M. luteus* (figure 1*b*). It is striking that the very similar Serosin-2 and Serosin-3 have such different antimicrobial specificity in this assay. One of the other upregulated effector genes, OFAS012870, is likely the previously described AMP Oncocin that is active against Gram negative human pathogens such as *Pseudomonas aeruginosa* and *Acinetobacter baumannii* [44]. Overall, our bacterial growth assay confirms the functionality of the innate immune reaction of *Oncopeltus* eggs.

**Figure 1.**
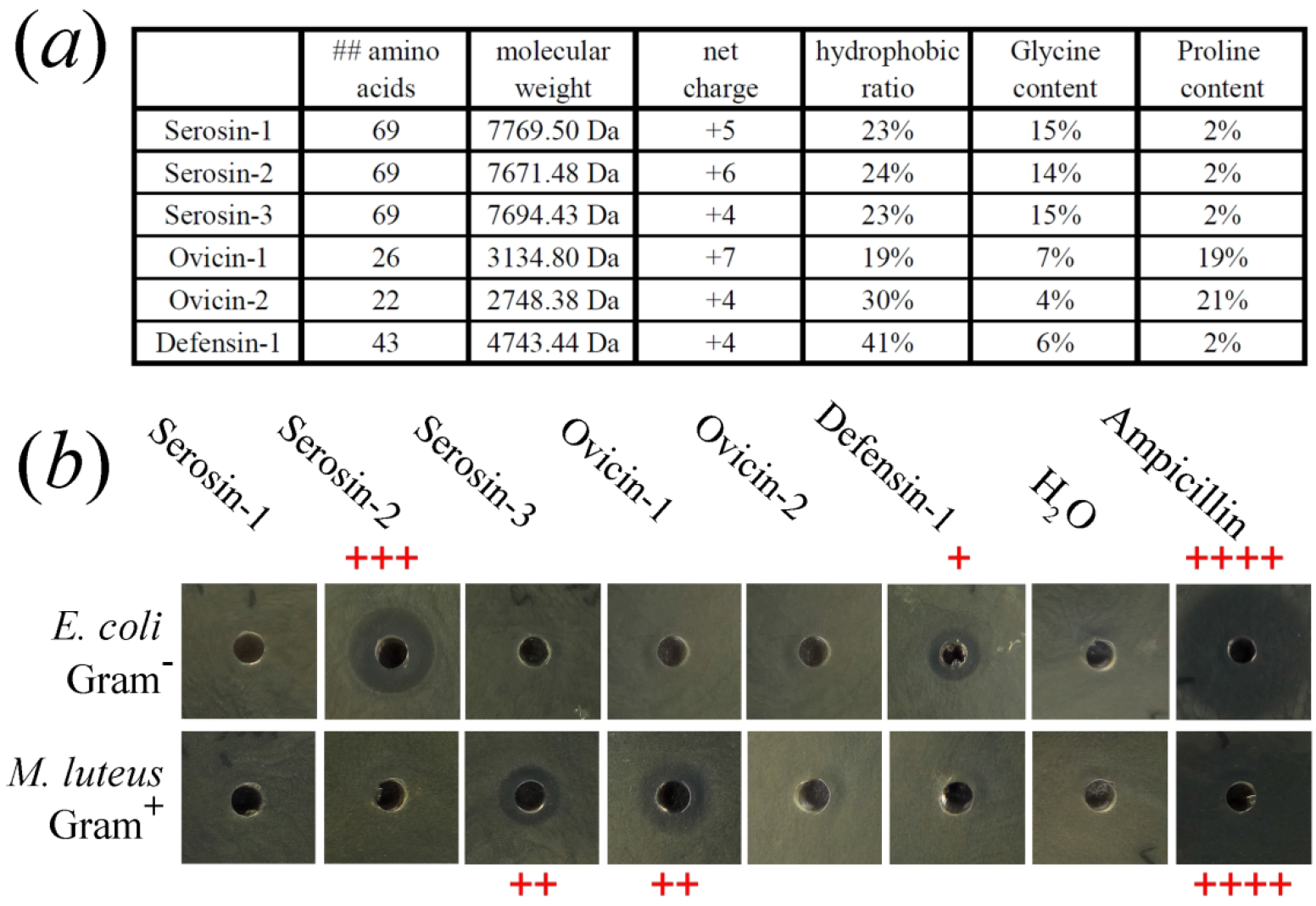
Analysis of novel potential antimicrobial *Oncopeltus* peptides. *(a)* Biochemical properties of Defensin-1 (OFAS005103) and five completely novel potential antimicrobial peptides predicted by software on the APD2 web site [42]. These peptides form two groups (called Serosins and Ovicins) of similar amino acid sequences encoded adjacently on the same strand of a chromosome. *(b)* Zone-of-inhibition assay. 1 nmole of synthesized peptide was applied to holes in an agar plate containing 1*10^5^ CFUs of either *Escherichia coli* (DMSZ 10514) or *Micrococcus luteus* (DMSZ 20030) per ml. Sterile water was used as negative control, and Ampicillin as positive control (1 mg/ml against *E. coli* and 100 μg/ml against *M. luteus*). Clear zones of inhibition were detected around Serosin-2 and Defensin-1 on plates containing Gram-negative bacteria, and around Serosin-3 and Ovicin-1 on plates containing Gram-positive bacteria.

To determine when *Oncopeltus* eggs gain this functional response, we performed qPCR for the six most strongly upregulated AMPs at three developmental stages 6 hours after sterile or septic injury. We chose stage A (16-24h) when the serosa has not yet developed; stage B (48-64h) when the serosa has enveloped the yolk and embryo; and stage C (112-120h) when the serosa has retracted during katatrepsis [5, 45] (figure 2, schematic drawings at the bottom). At the blastoderm stage, no transcriptional upregulation of AMPs was detected upon infection. In contrast, we detected strong upregulation of AMPs at stage B (figure 2). ANOVA analyses with post-hoc Tukey HSD reveal that upregulation after infection at stage B is significantly higher than at stage A, but not significantly different from stage C. This confirms that *Oncopeltus* eggs become immune responsive when the serosa has developed. But the eggs remain immune responsive after the serosa has disappeared (stage C, figure 2).

**Figure 2.**
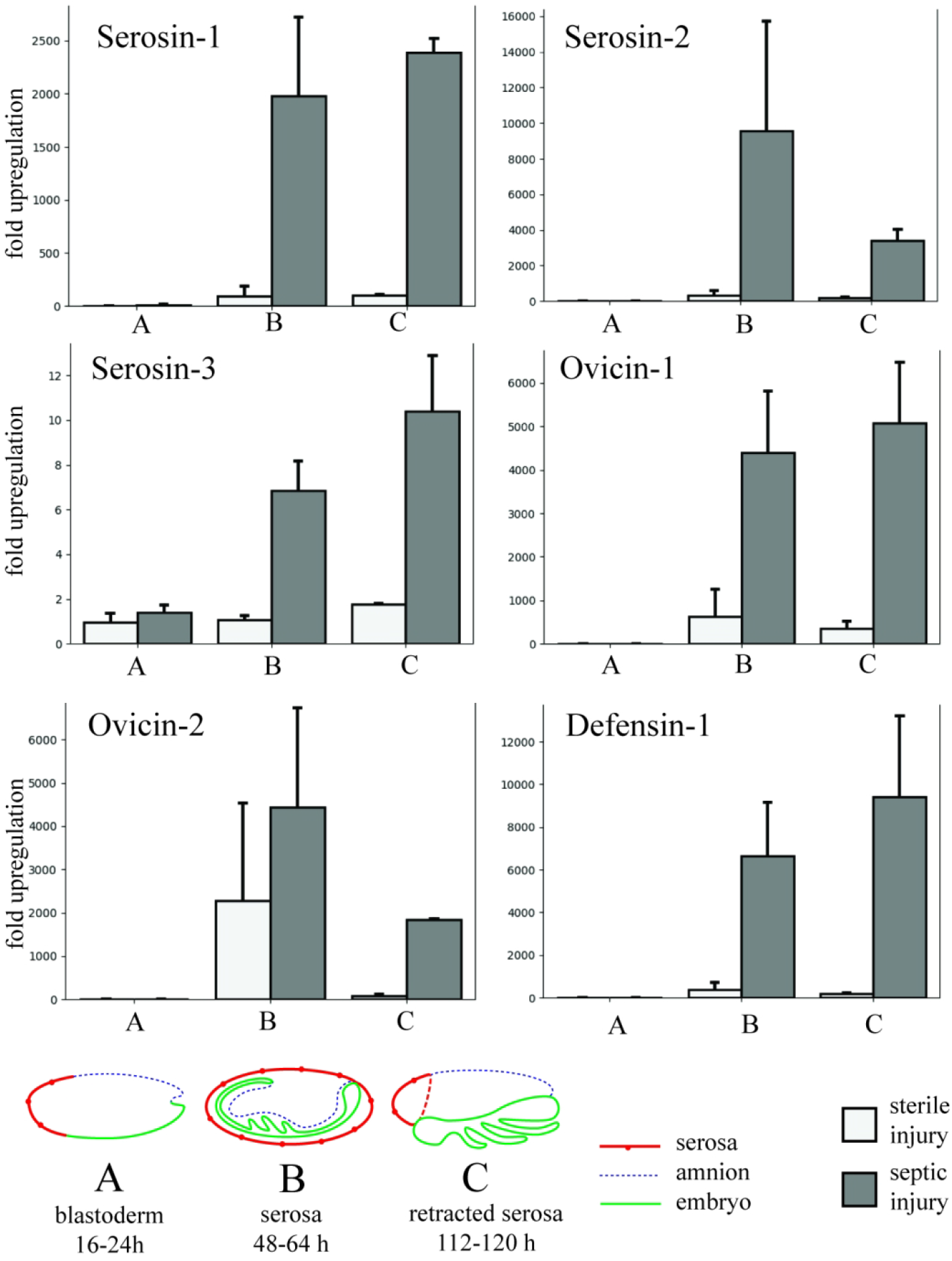
Upregulation of mRNAs for antimicrobial peptides after sterile or septic injury at three developmental stages of *Oncopeltus eggs*. Light grey bars: sterile injury. Dark grey bars: septic injury. Error bars show standard errors among three biological replicates (each biological replicate value is the mean of two technical replicate values). Stage A: blastoderm stage (16-24h old) when the serosa has not yet developed. Stage B: extending germ band stage (48-64h) when the serosa covers the developing embryo. Stage C: Retracted germ band stage (112-120 h) when the serosa has retracted after katatrepsis [5, 45]. Upon infection, upregulations of AMP genes at stage B and C are significantly higher than at stage A (p<0.001 for all genes, except for Serosin3 p<0.01), but not significantly different from each other (ANOVA with post-hoc Tukey HSD).

To reveal what tissue expresses the AMPs, we performed in situ hybridization for Serosin-1, Ovicin-2 and Defensin-1 at stages B and C after infection. We were unable to detect any expression of Ovicin-2 by in situ hybridization at any stage. In contrast, we found strong expression of Serosin and Defensin in the serosal cells at stage B, but not in the underlying embryo (figure 3*a-f*). This indicates that the serosa harbors the innate immune response of 48-64h old eggs. In stage C eggs, when the serosa has disappeared, we found expression of Serosin-1 in the developing mandibles and maxillae (figure 3*g, h*). Most conspicuously, expression was found in a row of patches in the dorsal ectoderm of the antennal, labial and thoracic segments, of which expression in the labial segment was strongest (figure 3*g*, dorsal view). The location of the expression is best demonstrated in a more ventral view of a different embryo (figure 3*h*). A very similar expression pattern was found for Defensin-1 (figure 3*j-l*). In addition, expression of Serosin-1 was found in the tracheae (figure 3*i*), the tissue that was found to be immune responsive in late *Drosophila* embryos too [22]. These expression patterns were not found after sterile injury or using a sense control in situ probe. We conclude that the serosa provides the *Oncopeltus* egg with an innate immune response until the ectoderm of the embryo proper becomes immune responsive.

**Figure 3.**
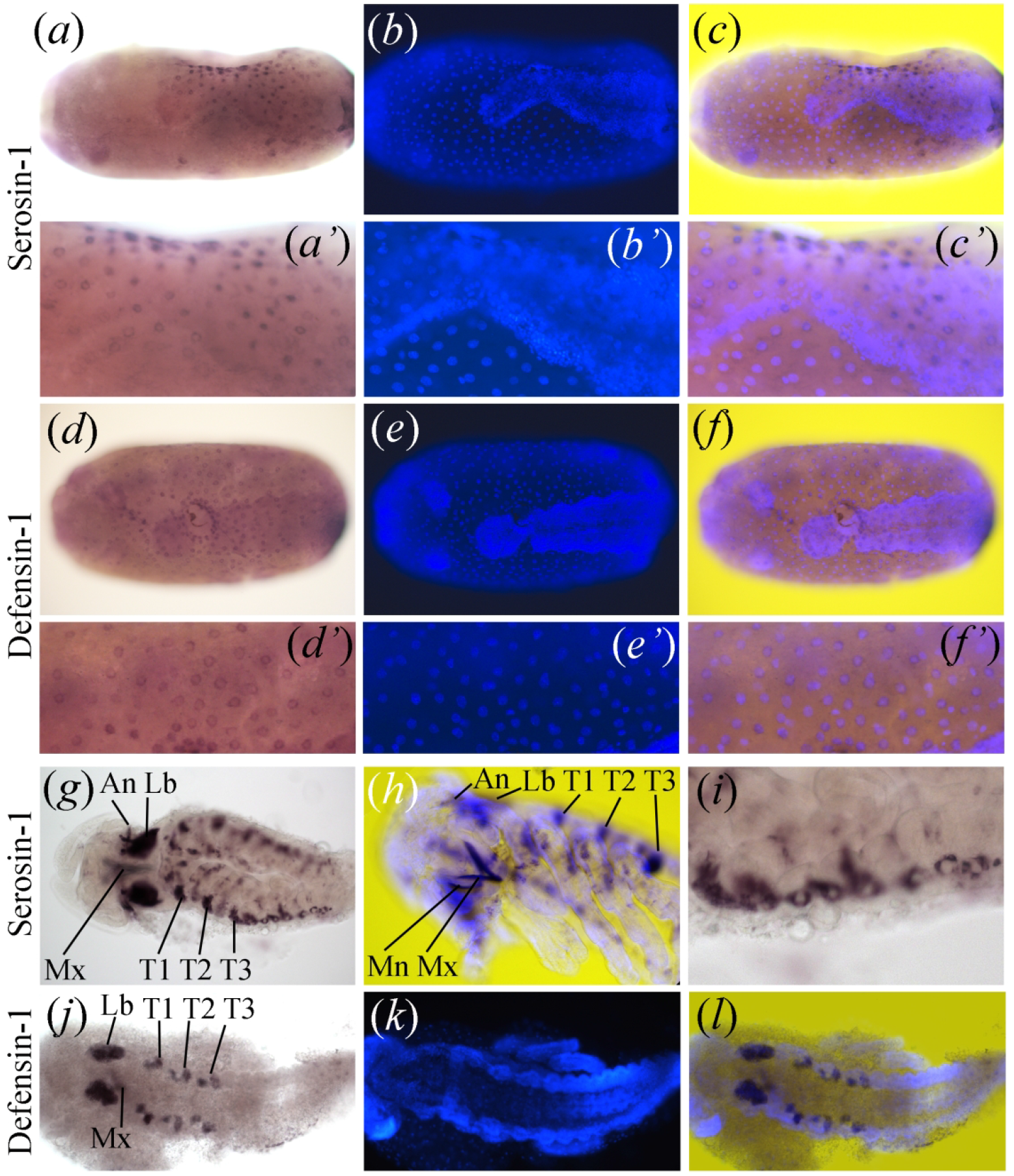
In situ hybridization for Serosin-1 and Defensin-1 six hours after immune challenge of *Oncopeltus* eggs. *(a-f)* Stage B embryos, when the serosa is present. *(g-l)* Stage C embryos, when the serosa retracts. (*a, d, g, i, j)* In situ signal (dark purple: colorimetric detection of an alkaline phosphatase-conjugated antibody bound to DIG-labelled probes, see materials and methods). *(b, e, k)* DAPI staining of the nuclei. *(c, f, h, l)* Overlay of in situ and DAPI staining (in situ layer has been permeabilized for the colour blue in Photoshop). *(a)* Serosin-1 is expressed around serosal nuclei around the septic injury (up). *(b)* DAPI counterstaining of *(a)*. The dense staining of the head and posterior of the germ band is visible under the serosal nuclei. *(c)* Overlay of *(a)* and *(b)*. *(a’)* Magnification of *(a)*. *(b’)* Magnification of B. *(c’)* Magnification of *(c)*. *(d)* Defensin-1 is expressed around the serosal nuclei upon septic injury. *(e)* DAPI counterstaining of *(d)*. The dense staining of the head and posterior of the germ band is visible under the serosal nuclei. *f)* Overlay of *(d)* and *(e)*. *(d’)* Magnification of *(d)*. *(e’)* Magnification of *(e)*. *(f’)* magnification of *(f)*. *(g)* Dorsal view. After the serosa has retracted, Serosin-1 is expressed in the maxillae, and in dorsal patches in the antennal, labial and thoracic segments. *(h)* Ventrolateral view. DAPI signal has been overlaid. Serosin-1 is expressed in mandibles and maxillae, and in dorsal patches in the antennal, labial, and thoracic segments. *(i)* After the serosa has retracted, Serosin-1 is expressed around the tracheae upon septic injury. *(j)* Dorsal view. After the serosa has retracted, Defensin-1 is detected in the maxillae, and in dorsal patches of the labial and thoracic segments. *(k)* DAPI counterstaining of *(j)*. *(l)* Overlay of *(j)* and *(k)*. Mx = maxillae, Mn = mandibles, An = antennal segment, Lb = labial segment, T1 = first thoracic segment, T2 = second thoracic segment, T3 = third thoracic segment.

Finally, we expanded our studies to the swarming grasshopper *Locusta migratoria* and evaluated immune competence of 72-84h old eggs by qPCR. At this stage, the serosa is present [46]. Our designed primers (see materials and methods) did not give any amplification in 10 cases (Locustin 2 and 3, Attacin1, 2 and 4, Defensin 2, 4, 5, 6, and putative defense protein 4), or unreliably high C_T_ values in all conditions (C_T_ > 37 for Attacin5, 6, 8 and 9, and Defensin 3). This could mean that these genes are not expressed in eggs, but could also indicate inefficient primer binding due to genetic variation, since our cDNA was derived from completely different locusts as was the genomic DNA used for genome assembly [47]. Nevertheless, for all AMPs for which we could detect expression, we observed increased expression upon infection, except for putative defense protein 1 (Figure 4*a-g*). Statistical analysis of the upregulation after the sterile and septic injury revealed significant upregulation upon infection of Locustin1, Attacin7 and Putative Defense Protein 2 (students t test as data were normally distributed) and defensin1 (Wilcoxon rank sum test as data were not normally distributed) (Figure4*a-g*). This shows that infection can induce expression of at least some antimicrobial peptides in the locust egg.

**Figure 4.**
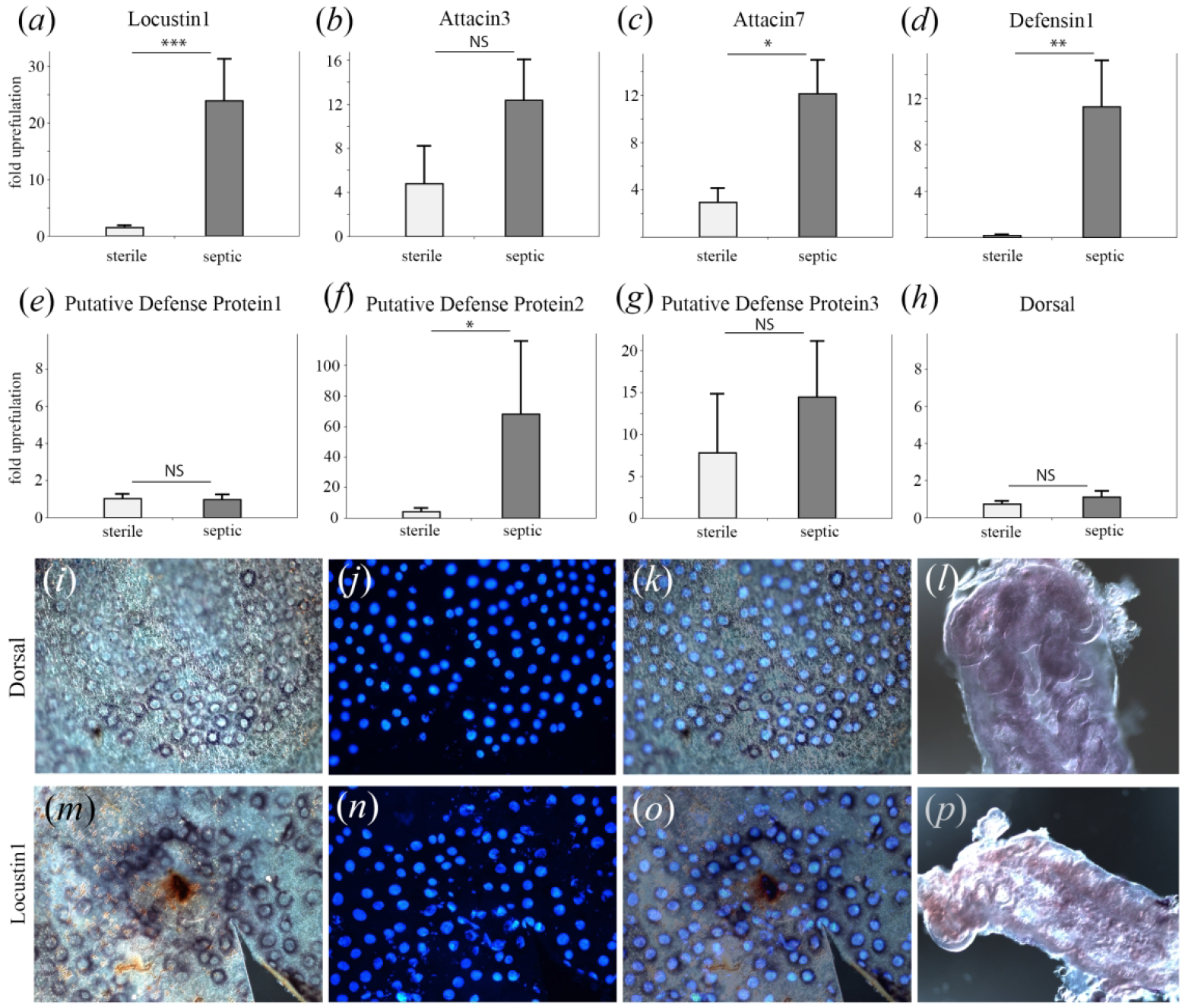
qPCR and in situ hybridization of immune gene expression in *Locusta migratoria* eggs. *(a-h)* Fold upregulation of immune genes 6h after sterile or septic injury of 72-84h old eggs compared to same-aged naïve eggs. Light grey bars: sterile injury. dark grey bars: septic injury. Error bars show standard errors among six biological replicates (each biological replicate value is the mean of two technical replicate values). Upregulation after septic injury compared to sterile injury in locust eggs is *(a)* highly significant for Locustin (p=0.0004, t=-5.24, df=10) *(b)* not significant for Attacin 3 (p=0.07, t=-2.01, df=10) *(c)* significant for Attacin7 (p=0.02, t=-2.76, df=10) *(d)* significant for Defensin1 (p=0.002, W=0) *(e)* not significant for Putative Defense Protein1 (p=0.82, t=0.24, df=10) *(f)* significant for Putative Defense Protein2 (p=0.01, t=-3.12, df=10) *(g)* not significant for Putative Defense Protein3 (p=0.07, t=-2.07, df=10) *(h)* not significant for Dorsal (p=0.391, t=-0.96097, df=4). *(i-l)* in situ hybridization for dorsal in unchallenged 72-84h old eggs. *(i)* The serosa shows constitutive expression of dorsal. *(j)* DAPI counterstaining of *(i)* shows the serosal nuclei. *(k)* Overlay of *(i)* and *(j)*. DAPI signal is quenched in a few instances by the in situ signal. *(l)* 72-84h old germ band does not show expression of dorsal. *(m-p)* In situ hybridization for Locustin1 6h after septic injury of 72-84h old eggs. *(m)* Strong expression of Locustin1 (dark purple) is visible in the serosa around the septic injury (brown melanization). *(n)* DAPI counterstaining of *(m)* shows the serosal nuclei. *(o)* Overlay of *(m)* and *(n)*. *(p)* The germband of challenged eggs does not show expression of Locustin1.

To investigate if the serosa is the immune competent tissue in locust eggs, we first performed in situ hybridization against the NFκB Dorsal, an important transcription factor in the innate immune response [48]. Dorsal is highly expressed in the serosa of *Tribolium*, which led to the first suggestion of an immune function for this extraembryonic epithelium [49]. In unchallenged *Locusta* eggs, dorsal is indeed expressed in the serosa (Fig.4*i-k*), and not in the 3 day old germ band of these eggs (figure 4*l*). As in *Tribolium*, dorsal mRNA is not significantly upregulated upon infection in the locust (figure 4*h*), as it is the mere nuclear translocation of the Dorsal protein that induces immune genes upon infection [48]. We could generate an in situ hybridization probe of sufficient length against the most significantly upregulated AMP, Locustin1, and observed no expression in unchallenged eggs, but clear expression around the septic injury in the serosa of challenged eggs (figure 4*m-o*). No specific staining was observed using dorsal or Locustin sense control probes. The constitutive expression of dorsal and induced expression of Locustin in the serosa strongly suggest that the serosa is the immune competent tissue in *Locusta migratoria* eggs.

Summarizing, qPCRs demonstrated an inducible immune response in eggs of the milkweed bug *Oncopeltus fasciatus* and the grasshopper *Locusta migratoria*. In situ hybridizations revealed that immune genes are not expressed in the young embryo, but in the serosa. We conclude that the serosa provides immune protection to these insect eggs until the ectoderm of the embryo itself becomes immune competent.

## 4. Discussion

We have provided the first evidence of an immune function for the serosal epithelium in eggs of two hemimetabolous insects. Together with the evidence from the holometabolous insects *Tribolium castaneum* and *Manduca sexta* [24, 25], this strongly suggests that immune competence is an ancestral feature of the serosa. Interestingly, immune competence of young eggs has been lost a few times in insect evolution. Eggs of the beetle *Nicrophorus vespilloides*, for instance, do not mount an innate immune response upon infection, although they do possess a serosa [17]. This might have been driven by a trade-off with developmental speed. *Nicrophorus* eggs develop within 2.5 days at 20°C which is exceptional among beetles [50]. Trade-offs between growth and immune competence are known from plants, birds and insects [51–53]. The loss of immune competence in *Nicrophorus* eggs might have been compensated by maternally provided lysozymes [54].

A similar trade-off with developmental speed might have taken place in *Drosophila. Drosophila melanogaster* lacks a serosa [12, 13] and early immune competence [21, 22]. Embryogenesis takes only 24 hours at 25°C, and the first instar larva already is immune competent shortly before hatching [55], probably greatly reducing the requirement of extensive egg protection. In *Ceratitis capitata*, another fruit fly without a serosa, maternal antimicrobial peptides have been found on the chorion of the eggs [56]. Thus, loss of immune competence in insect eggs may be compensated by maternal investments. Traditionally, studies on egg protection have focused on these parental investments (e.g. [56–58]). The well-studied social insects, for instance, provide extensive care for their eggs, protecting them against pathogens [59, 60]. In general, however, parental care is expensive and only arose in insects in the presence of intense selective pressures and specific behavioral precursors [61]. Indeed, recent evidence in the social termite rather suggests zygotic, endogenous protection, since antifungal activity of eggs arose from within the chorion and increased over developmental time [62]. Maternal mRNAs seem too short-lived to explain this increase of antimicrobial activity [63].

Along the same lines, enhanced survival of eggs in Transgenerational Immune Priming (TGIP) has traditionally been attributed to a maternal investment by loading of antimicrobials into the egg (e.g. [64–66]). In a comprehensive approach, such maternal loading was recently confirmed for *Tenebrio molitor* [67]. However, the role of a zygotic endogenous response in eggs is currently explicitly considered in TGIP studies (see [68, 69] for excellent reviews). Increasing antimicrobial activity over developmental time [70] and enhanced immune gene expression [71] upon parasitism in primed *Manduca sexta* eggs, for instance, suggest a zygotic response. Moreover, a series of papers has identified a mechanism that involves the binding of pathogen-associated molecules to vitellogenin [72], enabling the transfer of specific elicitors from the mother to the egg to induce a zygotic, endogenous response in these eggs [73–75].

The serosa is well suited to provide such zygotic, endogenous immune protection to the egg. The polyploid nuclei of the serosal cells allow the massive synthesis of peptides. The separation of the serosal cells from the embryo proper is the first visible differentiation step in the blastoderm of insects [76]. The serosa can thus protect the embryo long before the ectoderm starts to differentiate. The evolutionary origin of the serosa might have dramatically altered the costs for egg protection. Whereas crustaceans generally carry their eggs with them and make a large maternal investment [77], the serosal cells are of zygotic origin. Crustacean eggs have been reported to upregulate immune genes upon infection, but this upregulation seems to be rather weak (<4 fold upon infection maximally) and shortly before hatching [78]. The serosa has allowed insects to lay desiccation resistant and strongly immune competent eggs, likely relieving the need for parental care. Thus, the origin of the extra-embryonic serosa must have opened up a whole new range of oviposition sites for insects and may have contributed to their spectacular radiation as Pancrustacean group.

In conclusion, we have provided evidence of immune competence in the serosal epithelium of hemimetabolous insect eggs. This suggests that immune competence is an ancestral character of the serosa. The origin of the immune competent serosa might have greatly facilitated the success of the insects.

## Author-supplied statements

Relevant information will appear here if provided.

### Ethics

*Does your article include research that required ethical approval or permits?:*

This article does not present research with ethical considerations

*Statement (if applicable):*

#### CUST_IF_YES_ETHICS

No data available.

### Data

*It is a condition of publication that data, code and materials supporting your paper are made publicly available. Does your paper present new data?:*

Yes

*Statement (if applicable):*

The data discussed in this publication have been deposited in NCBIâ€™s Gene Expression Omnibus [33] and are accessible through GEO series accession number GSE100429 (https://www.ncbi.nlm.nih.gov/geo/query/acc.cgi?acc=GSE100429)

We used the following other datasets:

Murali, S. C., Bandaranaike, D., Bellair, M., Blankenburg, K., Chao, H., Dinh, H., Doddapaneni, H., Downs, B., Dugan-Rocha, S., Elkadiri, S., Gnanaolivu, R. D., Hernandez, B., Javaid, M., Jayaseelan, J. C., Lee, S., Li, M., Ming, W., Munidasa, M., Muniz, J., Nguyen, L., Ongeri, F., Osuji, N., Pu, L.-L., Puazo, M., Qu, C., Quiroz, J., Raj, R., Weissenberger, G., Xin, Y., Zou, X., Han, Y., Richards, S., Worley, K. C., Muzny, D. M., Gibbs, R. A., Koelzer, S. and Panfilio, K. A. (2015). Oncopeltus fasciatus genome assembly 1.0. Ag Data Commons. doi:10.15482/USDA.ADC/1173238

Vargas Jentzsch, Iris M.; Hughes, Daniel S.T.; Poelchau, Monica; Robertson, Hugh M.; Benoit, Joshua B.; Rosendale, Andrew J.; ArmisÃ©n, David; Duncan, Elizabeth J.; Vreede, Barbara M.I.; Jacobs, Chris G.C.; Berger, Chloe; Burnett, Denielle L.; Chang, Chun-che; Chen, Yen-Ta; Chipman, Ariel D.; Cridge, Andrew; CrumiÃre, Antonin; Dearden, Peter; Erezyilmaz, Deniz F.; Extavour, Cassandra; Friedrich, Markus; Horn, Thorsten; Hsiao, Yi-min; Jones, Jeffery W.; Jones, Tamsin E.; Khila, Abderrahman; Leask, Megan; Lovegrove, Mackenzie R.; Lu, Hsiao-ling; Lu, Yong; Nair, Ajay; Palli, Subba R.; Pick, Leslie; Porter, Megan L.; Refki, Peter N.; Rivera Pomar, Rolando; Roth, Siegfried; Sachs, Lena; Santos, Maria Emilia; Seibert, Jan; Sghaier, Essia; Shukla, Jayendra N.; Suzuki, Yuichiro; Tidswell, Olivia; Traverso, Lucila; van der Zee, Maurijn; Viala, Severine; Richards, Stephen; Panfilio, Kristen A.. (2020). Oncopeltus fasciatus Official Gene set v1.2. Ag Data Commons. https://doi.org/10.15482/USDA.ADC/1518752.

and the Locusta migratoria genome assembly L_migratoriav2 https://i5k.nal.usda.gov/bio_data/836515

### Conflict of interest

I/We declare we have no competing interests

*Statement (if applicable):*

#### CUST_STATE_CONFLICT

No data available.

### Authors’ contributions

This paper has multiple authors and our individual contributions were as below

*Statement (if applicable):*

C.G.C.J., R.v.d.H., Y.-T.C. R.P.W. and M.v.d.Z performed research. All authors analyzed data. C.G.C.J., R.v.d.H. and M.v.d.Z designed research. M.v.d.Z drafted the paper. C.G.C.J. and S.R. revised the paper

## Acknowledgements

We thank Tewodros Firdissa Duressa and Roger Huybrechts from KU Leuven for helping us to work with *Locusta migratoria*. We are grateful to Kees Koops for taking care of all insects. We thank the German Research Foundation (DFG) for funding (CRC 680). C.G.C.J. was supported by an Alexander von Humboldt postdoctoral fellowship. Y.-T. C. was supported by EMBO short term fellowship ASTF 362-2016.

## Availability of data and material

The data discussed in this publication have been deposited in NCBI’s Gene Expression Omnibus [33] and are accessible through GEO series accession number GSE100429 (https://www.ncbi.nlm.nih.gov/geo/query/acc.cgi?acc=GSE100429)

We used the following other datasets:

Murali, S. C., Bandaranaike, D., Bellair, M., Blankenburg, K., Chao, H., Dinh, H., Doddapaneni, H., Downs, B., Dugan-Rocha, S., Elkadiri, S., Gnanaolivu, R. D., Hernandez, B., Javaid, M., Jayaseelan, J. C., Lee, S., Li, M., Ming, W., Munidasa, M., Muniz, J., Nguyen, L., Ongeri, F., Osuji, N., Pu, L.-L., Puazo, M., Qu, C., Quiroz, J., Raj, R., Weissenberger, G.,

Xin, Y., Zou, X., Han, Y., Richards, S., Worley, K. C., Muzny, D. M., Gibbs, R. A., Koelzer, S. and Panfilio, K. A. (2015). Oncopeltus fasciatus genome assembly 1.0. *Ag Data Commons*.doi:10.15482/USDA.ADC/1173238

Vargas Jentzsch, Iris M.; Hughes, Daniel S.T.; Poelchau, Monica; Robertson, Hugh M.; Benoit, Joshua B.; Rosendale, Andrew J.; Armisén, David; Duncan, Elizabeth J.; Vreede, Barbara M.I.; Jacobs, Chris G.C.; Berger, Chloe; Burnett, Denielle L.; Chang, Chun-che; Chen, Yen-Ta; Chipman, Ariel D.; Cridge, Andrew; Crumière, Antonin; Dearden, Peter; Erezyilmaz, Deniz F.; Extavour, Cassandra; Friedrich, Markus; Horn, Thorsten; Hsiao, Yi-min; Jones, Jeffery W.; Jones, Tamsin E.; Khila, Abderrahman; Leask, Megan; Lovegrove, Mackenzie R.; Lu, Hsiao-ling; Lu, Yong; Nair, Ajay; Palli, Subba R.; Pick, Leslie; Porter, Megan L.; Refki, Peter N.; Rivera Pomar, Rolando; Roth, Siegfried; Sachs, Lena; Santos, Maria Emilia; Seibert, Jan; Sghaier, Essia; Shukla, Jayendra N.; Suzuki, Yuichiro; Tidswell, Olivia; Traverso, Lucila; van der Zee, Maurijn; Viala, Severine; Richards, Stephen; Panfilio, Kristen A.. (2020). Oncopeltus fasciatus Official Gene set v1.2. *Ag Data Commons*. https://doi.org/10.15482/USDA.ADC/1518752.

and the *Locusta migratoria* genome assembly L_migratoriav2 https://i5k.nal.usda.gov/bio_data/836515

## Authors’ contributions

C.G.C.J., R.v.d.H., Y.-T.C. R.P.W. and M.v.d.Z performed research. All authors analyzed data. C.G.C.J., R.v.d.H. and M.v.d.Z designed research. M.v.d.Z drafted the paper. C.G.C.J. and S.R. revised the paper. All authors read and approved the final manuscript

## SUPPLEMENTARY MATERIAL

Supplementary table 1

Manual annotation of all immune genes found in the *Oncopeltus fasciatus* genome [41] using our *de novo* assembly. OFASnumbers refer to *Oncopeltus fasciatus* Official Gene Set 1.2 [28]. CDS and protein sequences are included.

Supplementary table 2

Sequences of primers used for *Oncopeltus fasciatus* and *Locusta migratoria* qPCR upon sterile or septic injury.

Supplementary table 3

Mature peptide predictions for Serosins, Ovicins and Defensin-1 using SignalP [79] and manual corrections. OFASnumbers refer to *Oncopeltus fasciatus* Official Gene Set 1.2 [28].

Supplementary table 4

Significantly (adjusted p-value < 0.05) upregulated (>1.3 times) genes upon septic injury (DEseq output). OFASnumbers refer to *Oncopeltus fasciatus* Official Gene Set 1.2 [28].

Supplementary table 5

Significantly (adjusted p-value < 0.05) upregulated (>1.3 times) genes upon sterile injury (DEseq output). OFASnumbers refer to *Oncopeltus fasciatus* Official Gene Set 1.2 [28].

Supplementary table 6

Heatmap showing the number of RNA sequencing reads for all immune genes found in the *Oncopeltus fasciatus* genome (supplementary table 4). NA denotes Not Annotated and are mRNA models from a *de novo* assembly of our RNA sequencing reads. IMD, TAB and Toll9 are partial models and read counts may therefore be an underestimate

