## supplementary material, table 6 for "Immune function of the serosa in hemimetabolous insect eggs"

| Category | GeneID OFAS | Description | Naive | Sterile Injury | Septic Injury |
|---|---|---|---|---|---|
| Extracellular and Recognition | 014108 | Spaetzle | 69 | 109 | 98 |
| | 014107 | Spaetzle | 47 | 71 | 61 |
| | 002218 | Spaetzle | 39 | 35 | 38 |
| | 011266 | Spaetzle | | | |
| | 000511 | PGRP | 409 | 437 | 345 |
| | 001578 | PGRP | 57 | 179 | 195 |
| | 011838 | GNBP | 1263 | 1238 | 1218 |
| | 013163 | C-type lectin | 32 | 28 | 33 |
| | 010978 | C-type lectin | 2906 | 3552 | 3615 |
| | 017334 | C-type lectin | 12 | 19 | 16 |
| | 005206 | C-type lectin | 5298 | 4665 | 4838 |
| | 008425 | C-type lectin | 7425 | 7115 | 7429 |
| | 000862 | C-type lectin | 251 | 276 | 306 |
| | 010520 | Galectin | 29 | 40 | 38 |
| | 001654 | L-type lectin | 698 | 748 | 634 |
| | 001552 | L-type lectin | 2221 | 2188 | 2239 |
| | 014079 | FREP | 13 | 17 | 10 |
| | 009025 | Serine protease | 529 | 1804 | 2108 |
| | 006421 | Serine protease | 123 | 434 | 435 |
| | 009027 | Serine protease | 293 | 921 | 975 |
| | 006419 | Serine protease | 166 | 201 | 267 |
| Toll-pathway | 011521 | Toll1 | 10214 | [illegible] | [illegible] |
| | 002038 | Toll6 | 4000 | 3866 | 3697 |
| | 002798 | Toll7 | 3907 | 4125 | 3949 |
| | 005704 | Tollo / Toll8 | 5461 | 5477 | 5435 |
| | 009055 | Toll10 | 6644 | 6619 | 6904 |
| | 006184 | Toll9 | 8 | 5 | 11 |
| | 015942 | MyD88 | 336 | 377 | 318 |
| | 025126 | Tube-like kinase | 313 | 285 | 295 |
| | 025091 | Pelle | 1787 | 1905 | 1809 |
| | 014953 | Pellino | 1585 | 1614 | 1609 |
| | 002862 | TRAF6 | 473 | 419 | 449 |
| | 007860 | Cactus1 | 944 | 1707 | 1385 |
| | 012549 | Cactus2 - part1 | | | |
| | 018526 | Cactus2 - part2 | | | |
| | 016773 | Cactus3 | 53 | 53 | 59 |
| | 013817 | Cactus4 | 17 | 21 | 18 |
| | 002406 | Cactin | 1422 | 1423 | 1486 |
| | 007857 | Dorsal2 | 2531 | 2559 | 2580 |
| | 025092 | Dorsal | 508 | 846 | 601 |
| IMD-pathway | NA | Tab2 | 45 | 41 | 36 |
| | NA | FADD | 183 | 173 | 184 |
| | NA | IMD | 87 | 91 | 91 |
| | 017036 | Dredd | 1889 | 1829 | 1825 |
| | 003620 | IAP2 | 1452 | 1544 | 1391 |
| | 005964 | TAK1 | 309 | 317 | 304 |
| | 006790 | Caspar | 3793 | 4006 | 3786 |
| | 011587 | IKK epsilon | 239 | 214 | 212 |
| | 018147 | Relish - part1 | 223 | 207 | 221 |
| | 013061 | Relish - part2 | 784 | 727 | 813 |

| | | |
|---|---|---|
| **JNK-pathway** | 001126 | Hemipterous |
| | 016451 | JNK |
| | 002923 | Jra |
| | 019014 | Kayak |

| | | |
|---|---|---|
| **JAK-STAT pathway** | 025220 | Domeless |
| | 002331 | Hopscotch |
| | 019212 | STAT 5B-like - part1 |
| | 010550 | STAT 5B-like - part2 |
| | 009084 | JAK |

| | | |
|---|---|---|
| **Effectors** | 005107 | Defensin |
| | 005103 | Defensin |
| | 005106 | Defensin |
| | 005105 | Defensin |
| | 005104 | Defensin |
| | 008569 | Defensin |
| | 008940 | Defensin |
| | 008943 | Defensin |
| | 008939 | Defensin |
| | 011278 | Defensin |
| | 011277 | Defensin |
| | 008937 | Defensin |
| | 008942 | Defensin |
| | 008938 | Defensin |
| | 008941 | Defensin |
| | 017707 | Hemiptericin |
| | 017709 | Hemiptericin |
| | 017708 | Hemiptericin |
| | 018152 | Hemiptericin |
| | 018151 | Hemiptericin |
| | 012870 | Oncocin |
| | 004519 | Potential AMP |
| | 018039 | Potential AMP |
| | 009172 | Potential AMP |
| | 006190 | Potential AMP |
| | 003658 | Potential AMP |
| | 013043 | Potential AMP |
| | 009533 | Potential AMP |
| | 009139 | Ovicin1 |
| | 009140 | Ovicin2 |
| | 013883 | Serosin1 |
| | 013884 | Serosin2 |
| | 013885 | Serosin3 |
| | 014551 | Hexamerin |
| | 013226 | C-type lysozyme |
| | 015457 | I-type lysozyme |
| | 015456 | I-type lysozyme |
| | 015454 | I-type lysozyme |
| | 009274 | I-type lysozyme |
| | 015455 | I-type lysozyme |
| | 009275 | I-type lysozyme |
| | 000034 | prophenoloxidase |
| | 018384 | prophenoloxidase |

no reads
< 10
11-50
51-100
101-200
201-300
301-400
401-500
501-750
751-1000
1001-1250
1251-1500
1501-2500
2501-10000
10001-12500
12500+
